# The evolution of sex differences in disease genetics

**DOI:** 10.1101/000414

**Authors:** William P. Gilks, Jessica K. Abbott, Edward H. Morrow

## Abstract

There are significant differences in the biology of males and females, ranging from biochemical pathways to behavioural responses, which are relevant to modern medicine. Broad-sense heritability estimates differ between the sexes for many common medical disorders, indicating that genetic architecture can be sex-dependent. Recent genome-wide association studies (GWAS) have successfully identified sex-specific and sex-biased effects, where in addition to sex-specific effects on gene expression, twenty-two medical traits have sex-specific or sex-biased loci. Sex-specific genetic architecture of complex traits is also extensively documented in model organisms using genome-wide linkage or association mapping, and in gene disruption studies. The evolutionary origins of sex-specific genetic architecture and sexual dimorphism lie in the fact that males and females share most of their genetic variation yet experience different selection pressures. At the extreme is sexual antagonism, where selection on an allele acts in opposite directions between the sexes. Sexual antagonism has been repeatedly identified via a number of experimental methods in a range of different taxa. Although the molecular basis remains to be identified, mathematical models predict the maintenance of deleterious variants that experience selection in a sex-dependent manner. There are multiple mechanisms by which sexual antagonism and alleles under sex-differential selection could contribute toward the genetics of common, complex disorders. The evidence we review clearly indicates that further research into sex-dependent selection and the sex-specific genetic architecture of diseases would be rewarding. This would be aided by studies of laboratory and wild animal populations, and by modelling sex-specific effects in genome-wide association data with joint, gene-by-sex interaction tests. We predict that even sexually monomorphic diseases may harbour cryptic sex-specific genetic architecture. Furthermore, empirical evidence suggests that investigating sex-dependent epistasis may be especially rewarding. Finally, the prevalent nature of sex-specific genetic architecture in disease offers scope for the development of more effective, sex-specific therapies.

**Funding:** This work was supported by the European Research Council (WPG and EHM; Starting Grant #280632), a Royal Society University Research Fellowship (EHM), the Swedish Research Council (JKA), and the Volkswagen Foundation (JKA). The funders had no role in study design, data collection and analysis, decision to publish, or preparation of the manuscript.

**Competing interests:** The authors declare that they have no competing financial interests.

## 1. Introduction

Sex, and the existence of two sexes, has revolutionised life on Earth. The success of sexual reproduction is attributed to recombination between parental chromosomes, which accelerates the loss of deleterious alleles and the proliferation of advantageous ones [1,2]. The difference in gamete size between males and females is a fundamental property of almost all sexual species. Yet sexual dimorphism extends far beyond this, from cellular and anatomical specialisation to secondary sexual traits such as ornamentation and behaviour. Furthermore, there are differences in gene co-expression and metabolome networks between the sexes [3–5]. It is therefore not surprising that in the field of medicine, males and females frequently differ in core features of disease [6].

The genetic basis of disease has been intensely researched, with the aim of providing improved diagnosis and therapy. Heritable diseases can be classified as being rare with monogenic aetiology (caused by a single mutation), or common (prevalence 0.1-1%), caused by multiple genetic variants, each with high population frequency but small individual contribution to disease risk [7,8]. For these genetically complex diseases and traits, genome-wide association studies (GWAS) have been successful at identifying loci, but the heritability accounted for by main effects, and by polygenic risk score, remains conspicuously low [9,10]. This deficit is stimulating the consideration of other factors such as the environment and epistasis [11]. Sex differences in the genetic architecture of common diseases have been known for some time [12], and recent analysis of large GWAS datasets has resulted in an unprecedented rise in the number of known sex-specific loci for human diseases and quantitative traits (see next section). Whilst this fact alone should encourage further investigation, evolutionary theory also predicts the existence of sex-specific genetic architecture for complex traits via sex-specific or sexually antagonistic selection.

Males and females share genes and genetic variation (excepting the Y chromosome), yet often have divergent optimum conditions for survival and reproduction [13]. A recent model predicts that even a marginal difference in selection pressure rapidly amplifies the contribution deleterious alleles make to trait architecture [14]. Additionally, opposing selection pressures on a shared trait creates sexual antagonism, in which the strength of positive selection for an allele in one sex will allow it to be maintained even if it is deleterious to the other sex (see section 3) [15,16]. In this review we summarise recent evidence for the sex-specific genetic architecture of common diseases, and the evidence for sexual antagonism in the light of evolutionary theory. We also propose new mechanisms by which sexual antagonism can contribute towards the genetic architecture of disease, and guidelines for the identification of sex-specific genetic effects.

## 2. Evidence for sex-specific genetic architecture

Broad-sense heritability is the proportion of phenotypic variance in a population sample that can be attributed to genetic variation [17]. With identical genetic architecture, and assuming a common environment, trait heritability should be equal in male and female samples. However, in a study of twenty quantitative traits in humans, eleven showed significant sex-bias in heritability [18]. Following a PubMed literature search, we identified eighteen independent studies in humans (representing thirty-one traits) that provided separate heritability estimates for males and females, and also stated whether the difference was statistically significant. A summary of these data is presented in Figure 1, showing that while fifteen traits did not exhibit significant sex-bias in heritability, thirteen had a higher heritability in females, and three a higher heritability in males. Although there may be some bias in these studies (non-reporting, non-independence of traits or prior selection of traits with known sexual dimorphism), they illustrate that the heritability of complex traits is commonly sex-biased across a range of phenotype classes.

**Figure 1:**
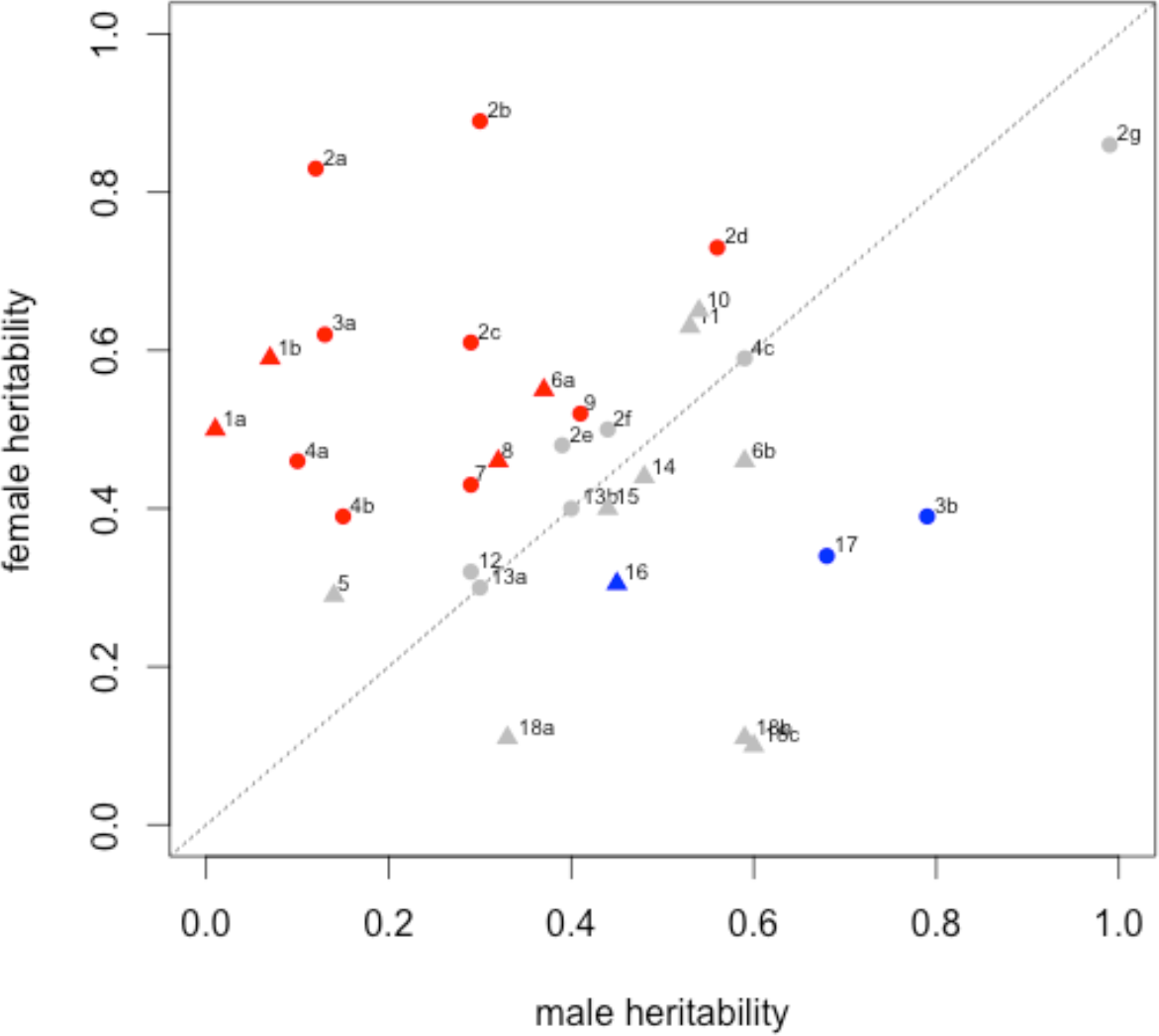
Comparison of male vs female narrow-sense heritability estimates from human studies. Red and blue-coloured data points indicate significant differences between the sexes. Grey data points indictate no significant difference between the sexes. Triangles indicate a binary/qualitative phenotype. Circles indicate a continuous/quantitative phenotype. 1a: Drive for thinness. 1b Body Dissatisfaction [103]. 2a: Waist diameter. 2b: Waist-height ratio. 2c: Body-mass index. 2d: Peripheral body fat. 2e: Hip diameter. 2f: Body weight. 2g: Body height [104]. 3a: Triglyceride serum level. 3b: LDL cholesterol serum level [105]. 4a: Lung FEV1 (forced exit volume). 4b: Lung D_LCO_ (diffusing capacity). 4c: Lung VC (vital capacity) [106]. 5: Geriatric depression [107]. 6a: Smoking initiation. 6b: Regular tobacco use [108]. 7: Sleep reactivity (insomnia) [109]. 8: Alcohol dependence [110]. 9: Subjective well-being [111]. 10: Reading disability [112]. 11: Reading difficulties [113]. 12: Self-esteem [114]. 13a: Respiratory sinus arythmia. 13b: Heart beat entropy [115]. 14: Tension-type headache [116]. 15: Lower back pain [117]. 16: Seasonal mood change [118]. 17: Protein C sensitivity [119]. 18a: Drug use. 18b: Tobacco use. 18c: Alcohol use [120].

It is well known that non-genetic factors influence differences in heritability, the most obvious being sex hormones (androgens, oestrogens and progesterones, secreted from the gonads). These can create systemic differences between males and females for trait expression, which in turn affects disease risk and heritability, for example the protective effect of oestrogen on heart disease [19]. However, experiments using hormone treatment and gonadectomy show that some gender differences in phenotypes, such as immune response, behaviour, and toxin resistance, are not determined by sex hormones but by sex chromosome dosage [20–22]. This implies that heritability differences are not always caused by sex hormones, and can be caused by sex-specific differences in genetic architecture, whereby a genetic variant has a different phenotypic outcome depending on whether it is expressed in a male or female environment.

The molecular genetic evidence for sex-specific genetic architecture is strong. For gene expression in human cell lines, 15% of SNPs that control gene expression (expression quantitative trait loci or eQTL) do so in a sex-specific manner, even in the absence of sex hormones [23]. For complex traits, GWAS have identified many robust sex-specific loci across a range of human phenotypes. These results are summarised Table 1, which shows thirty-two loci with sex-dependent effects in the twenty-two traits studied. The majority of the effects were sex-specific (twenty-eight loci significant in one sex only) although five sex-biased effects were also reported (significant in both sexes but different magnitude of effect). One opposite effect direction locus has also been reported from a GWAS (for recombination rate [24]). Model organisms such as laboratory mice *Mus musculus* and the fruit fly *Drosophila melanogaster* have been used successfully in many phenotype-mapping projects, facilitated by the controlled environment and flexibility of experimental design. Indeed, many sex-specific eQTL have been identified in the fruit fly *D. melanogaster* [25] and laboratory mice [26]. For genetically complex traits, sex-specific loci have been identified for sleep patterns and ageing in *D. melanogaster*, and for fat mass and skeletal traits in mice [27–30].

**Table 1:** Loci with sex-dependent effects on human phenotypes, identified through genome-wide SNP or CNV analysis

| Phenotype | Sex test | Individuals tested | Gene | Chromosome band | SNP | MAF | Male effect<sup>†</sup> | Female effect<sup>†</sup> | Reference |
| --- | --- | --- | --- | --- | --- | --- | --- | --- | --- |
| Mitochondrial DNA levels | Separate | 384 | MRPL37 | 1p32.3 | rs10888838 | 0.11 | 0.81 | ns | [121] |
| Heart beat rate (QT interval) | Separate | 3761 | NOS1AP | 1q23.3 | rs10494366 | 0.29 | 3.08 | 2.09 | [122] |
| Waist-height ratio | Sep & p-diff | 175585 | LYPLAL1/SLC30A10 | 1q41 | rs4846567 | 0.29 | ns | 0.06 | [123] |
| Waist-height ratio | Separate | 190803 | " | " | rs2820443 | 0.29 | ns | 0.05 | [124] |
| Visceral adiposity | Separate | 117857 | THNSL2 | 2p11.2 | rs1659258 | 0.35 | ns | Z-score 1.5 | [125] |
| Mitochondrial DNA levels | Separate | 384 | RNF144 | 2p25.1 | rs2140855 | 0.39 | ns | 0.32 | [121] |
| Waist-height ratio | Sep & p-diff | 175585 | GRB14/COBLL1 | 2q24.3 | rs10195252 | 0.44 | ns | 0.05 | [123] |
| " | Separate | 190803 | " | " | rs6717858 | 0.44 | ns | 0.05 | [124] |
| Plasma homocysteine | Separate | 1679 | CPS1 | 2q34 | rs1047891 | 0.30 | ns | 0.04 | [126] |
| Glycine levels | Separate | 3343 | " | " | rs715 | 0.24 | ns | 0.23 | [3] |
| Crohn's Disease | Separate | 8463 | ATG16L1 | 2q37.1 | rs3792106 | 0.40 | ns | OR 1.48 | [127] |
| Waist-height ratio | Sep & p-diff | 175585 | PPARG | 3p25.2 | rs4684854 | 0.42 | ns | 0.04 | [124] |
| Waist-height ratio | Sep & p-diff | 175585 | ADAMTS9 | 3p14.1 | rs6795735 | 0.19 | ns | 0.05 | [123] |
| Recombination rate | Separate | 2500 | RNF212 | 4p16.3 | rs11939380 | 0.33 | +118cM | ns | [128] |
| " | Separate | 19578 | " | " | rs1670533 | 0.22 | -69.4cM | +88.2cM | [24] |
| Uric acid concentration | p-diff | 28141 | SLC2A9 | 4p16.1 | rs734553 | 0.26 | -0.22 | -0.40 | [129] |
| Sex-hormone binding globulin | Separate | 21791 | UGT2B15 | 4q13.2 | rs293428 | 0.30 | -0.03 | ns | [130] |
| Uric acid concentration | p-diff | 28141 | ABCG2 | 4q22.1 | rs2231142 | 0.12 | 0.22 | 0.13 | [129] |
| Waist circumference | Sep & p-diff | 199499 | MAP3K1 | 5q11.2 | rs11743303 | 0.19 | ns | 0.03 | [124] |
| Low-density lipoprotein (LDL) | Sep & p-diff | 20512 | HMGCR | 5q13.3 | rs12654264 | 0.38 | -4.03 | ns | [131] |
| Thyroid stimulating hormone | Separate | 26420 | PDE8B | 5q13.3 | rs6885099 | 0.29 | -0.17 | -0.12 | [132] |
| Naso-pharyngeal cancer | Separate | 1437 | MICA/HCP5 | 6p21.33 | na (CNV) | 0.03 | OR 3-141 | ns | [133] |
| Waist-height ratio | Sep & p-diff | 175585 | VEGFA | 6p21.1 | rs6905288 | 0.45 | ns | 0.05 | [123] |
| Thyroid stimulating hormone | Separate | 26420 | PDE10A | 6q27 | rs753760 | 0.50 | 0.13 | 0.08 | [132] |
| Pro-insulin levels | GWAMA* | 27079 | DDX31 | 9q34.13 | rs306549 | 0.24 | 0.04 | ns | [88] |
| Obesity & Osteoporosis (bivariate) | Separate | 4355 | SOX6 | 11p15.1 | rs297325 | 0.20 | OR 1.75 | ns | [134] |
| Triglyceride levels | Sep & p-diff | 24273 | APOA5/BUD13 | 11q23.3 | rs28927680 | 0.07 | 0.13 | ns | [131] |
| Type II Diabetes | Separate | 149000 | CCND2 | 12p13.32 | rs11063069 | 0.21 | OR 1.08-1.16 | ns | [135] |
| Type I Diabetes<sup>‡</sup> | Separate | 27530 | CTSH | 15q25.1 | rs3825932 | 0.30 | OR 1.13-1.27 | ns | [136] |
| Thyroxine levels (FT4) | Separate | 17498 | LPCAT2/CAPNS2 | 16q12.2 | rs6499766 | 0.48 | 0.02 | ns | [132] |
| Thyroid stimulating hormone | Separate | 26420 | MAF | 16q23.2 | rs3813582 | 0.38 | 0.12 | 0.06 | [132] |
| Recombination rate | Separate | 2500 | 17q21.31 region | 17q21.31 | rs2668622 | 0.20 | ns | +124cM | [128] |
| Height | Separate | 1625 | NEDD4L | 18q21.31 | CNP12587 | 0.02 | ns | -8.1% | [137] |
| Thyroxine levels (FT4) | Separate | 17146 | NETO1/FBXO15 | 18q22.3 | rs7240777 | 0.47 | ns | -0.08 | [132] |
| Type II Diabetes | Separate | 149000 | GIPR | 19q13.32 | rs8108269 | 0.31 | ns | OR 1.06-1.14 | [135] |
| High-density lipoprotein (HDL) | Sep & p-diff | 11528 | PLTP | 20q13.12 | rs7679 | 0.18 | ns | 1.68 | [131] |

MAF Minor allele frequency. Value for similar HapMap population sample stated when study sample MAF not available
<sup>†</sup> Effect value is for the correlation coefficient $\beta$ unless otherwise stated. OR Odds ratio, 95% confidence intervals.
<sup>‡</sup> Result of separate-sex analysis of SNPs previously identified in a standard, main-effects analysis.
\* GWAMA 'Genome-wide analysis, meta-analysis' (Magi et al 2010)
SNP rs1047891 previously known as rs7422339.

Gene manipulation studies of model organisms have identified hundreds of genes with sexually dimorphic effects on disease-related phenotypes. Interestingly, not only have sex-specific and sex-biased effects been observed, but also sex-reversed and sexually pleiotropic effects (when the same locus affects different traits in males and females; see section 5). For example, murine vitamin D receptor disruption causes weight loss in males but decreased bone density in females [31], and p53 over-expression in *D. melanogaster* increases male life-span but reduces that of females [32]. Furthermore, a screen of 1,332 *D. melanogaster* P-element insertion lines identified forty-one mutations that had sexually dimorphic effects on life-span including six that were in opposite directions [33]. Although gene disruption studies do not precisely reproduce the effect of natural genetic variation, they demonstrate that different pathways can control the same trait. The question then is: how does this sexual dimorphism in genetic effects arise? Insights from evolutionary biology are of great value here, since theory about the origin and evolution of sex differences is well-developed, both on the phenotypic and on the genetic level.

## 3. Sexual antagonism

Sexual antagonism is a genetic conflict resulting from sex-specific selection acting on a shared genome. A subcategory of sexual antagonism is intra-locus sexual conflict, where a trait is controlled by the same genes, and is distinct from inter-locus sexual conflict, which concerns reproductive interactions involving different loci in each sex. Sexual antagonism has now been demonstrated in a wide variety of taxa, including plants, birds, mammals, and insects [16,34]. Anisogamy (difference in gamete size) is considered to be the ultimate source of sex-specific selection [35,36], although ecological factors can also play a role in shaping patterns of sex-specific selection [37]. The fact that males produce many small gametes and females few large gametes means that reproductive strategies are fundamentally different across the sexes, which is thought to result in the evolution of sexual dimorphism [38]. However these divergent phenotypes must be developed from a common pool of genetic information, making it difficult to simultaneously achieve optimum trait values in both sexes. Thus, for certain traits a conflict will be maintained and the sexes will be displaced from their optimum phenotypes. For example, when selection on females was completely removed, experimentally evolved fruit flies became masculinized in a number of traits demonstrating that males had previously been displaced from their phenotypic optimum by counter-selection in females (reviewed in [39]). Pedigree analysis of wild animal populations has also demonstrated a negative intersexual genetic correlation for fitness i.e. genotypes producing successful males produce unsuccessful females and vice versa [40,41].

More formally, sexual antagonism occurs when genetically correlated traits have opposite effects on male and female fitness. In the simplest case, increasing values of a single trait would increase fitness in one sex and decrease it symmetrically in the other sex (Figure 2a). In this case the trait is assumed to be positively correlated between the sexes. However more complicated patterns are also possible, such as opposite fitness effects of different correlated traits (Figure 2b-c) or asymmetric patterns of selection (Figure 2d). Consistent with this, a recent study demonstrated that human height was likely to be subject to sexual antagonism: within sibling pairs, men of average height had higher fitness while shorter women had higher fitness [42]. This means that the fitness effect of a given height-determining allele will be context-dependent in terms of sex, and that the population as a whole will be unlikely to evolve towards a shorter phenotype, despite directional selection in females, because of counter-selection in males. One of the major evolutionary implications of sexual antagonism is the maintenance of genetic variation that is deleterious to one sex. Although this has not been fully demonstrated at the molecular level, the population dynamics of a synthetic sexually antagonistic allele in a laboratory *D. melanogaster* study accurately follows predictions of mathematical models [43,44].

**Figure 2:**
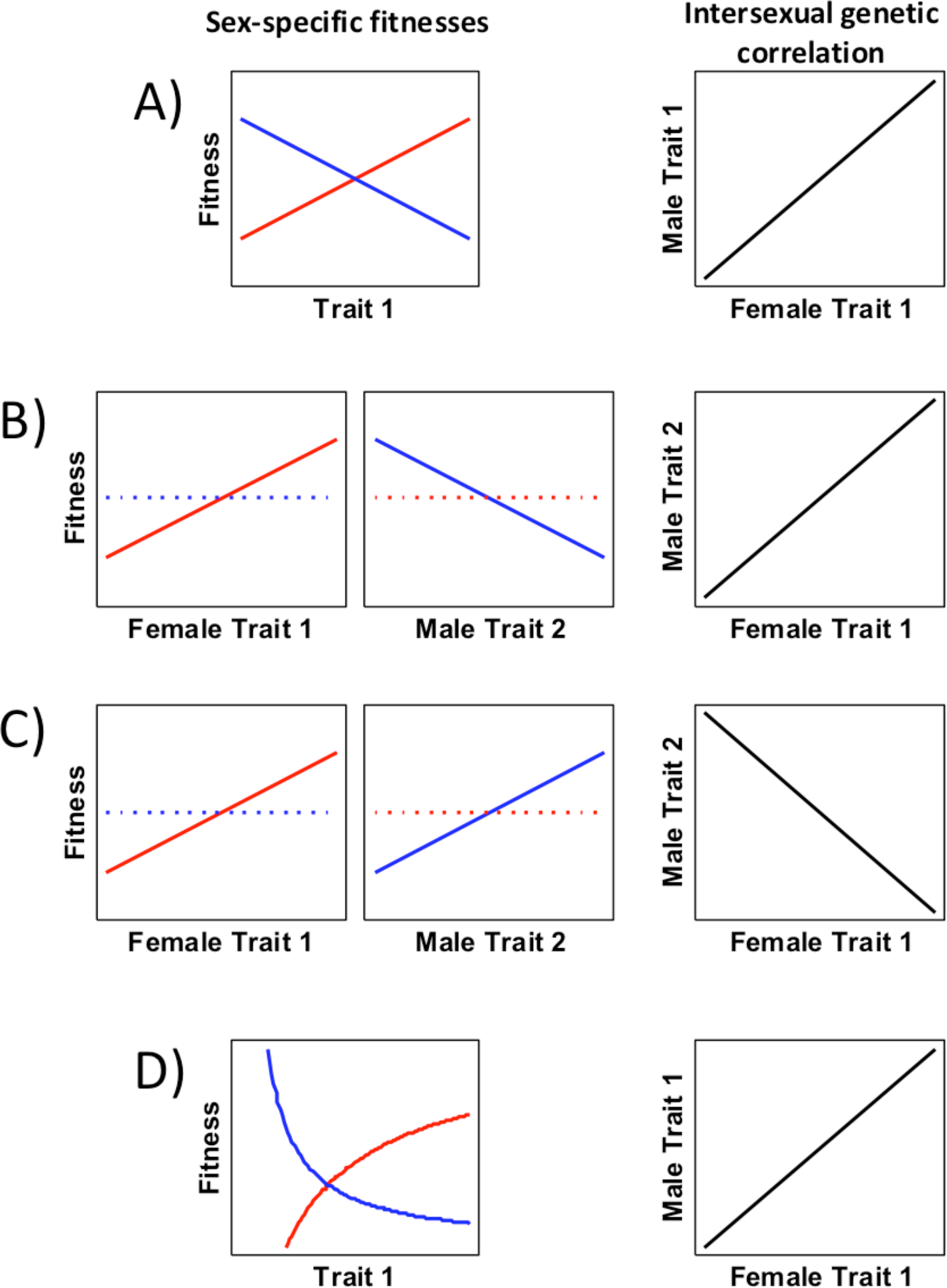
The various forms sexual antagonism can take. Female fitness functions are shown with red lines, male with blue lines, and the intersexual genetic correlation with black lines. A. The simplest case (also known as intralocus sexual conflict) is where the same trait has opposite and approximately symmetric fitness effects on males and females. The intersexual genetic correlation for the traits is high and positive. B. Sexual antagonism can also occur when different traits have a high positive intersexual genetic correlation, but are selected in opposite directions in males relative to females. In the unselected sex (broken lines), selection for the trait in question might be weakly positive, neutral, or even absent if the trait is sex-limited. C. Although no empirical examples of this type have yet been demonstrated, it is also possible that traits with a strong negative intersexual genetic correlation could be subject to sexual antagonism, assuming both traits are selected concordantly across the sexes. A negative intersexual genetic correlation could occur when the same gene product is incorporated in competing alternative pathways. D. It should also be pointed out that selection pressures need not be completely symmetric. Non-linear relationships are also possible.

## 4. Sexual dimorphism and resolution of sexual antagonism

A sexually antagonistic trait is expected to go through several evolutionary stages (see Figure 3 for more detail [16,38]).

1. Initially, the trait is monomorphic and under weak stabilizing selection.
2. A change in the physical or social environment causes the previously concordant trait to become subject to opposite patterns of sex-specific directional selection. This is the most severe stage of antagonism.
3. Sexual dimorphism then evolves, causing the sexes to come closer to their respective phenotypic optima, but the antagonism is only partially resolved.
4. Finally, the sexes reach their optimal phenotype values and the sexual antagonism is completely resolved.

**Figure 3:**
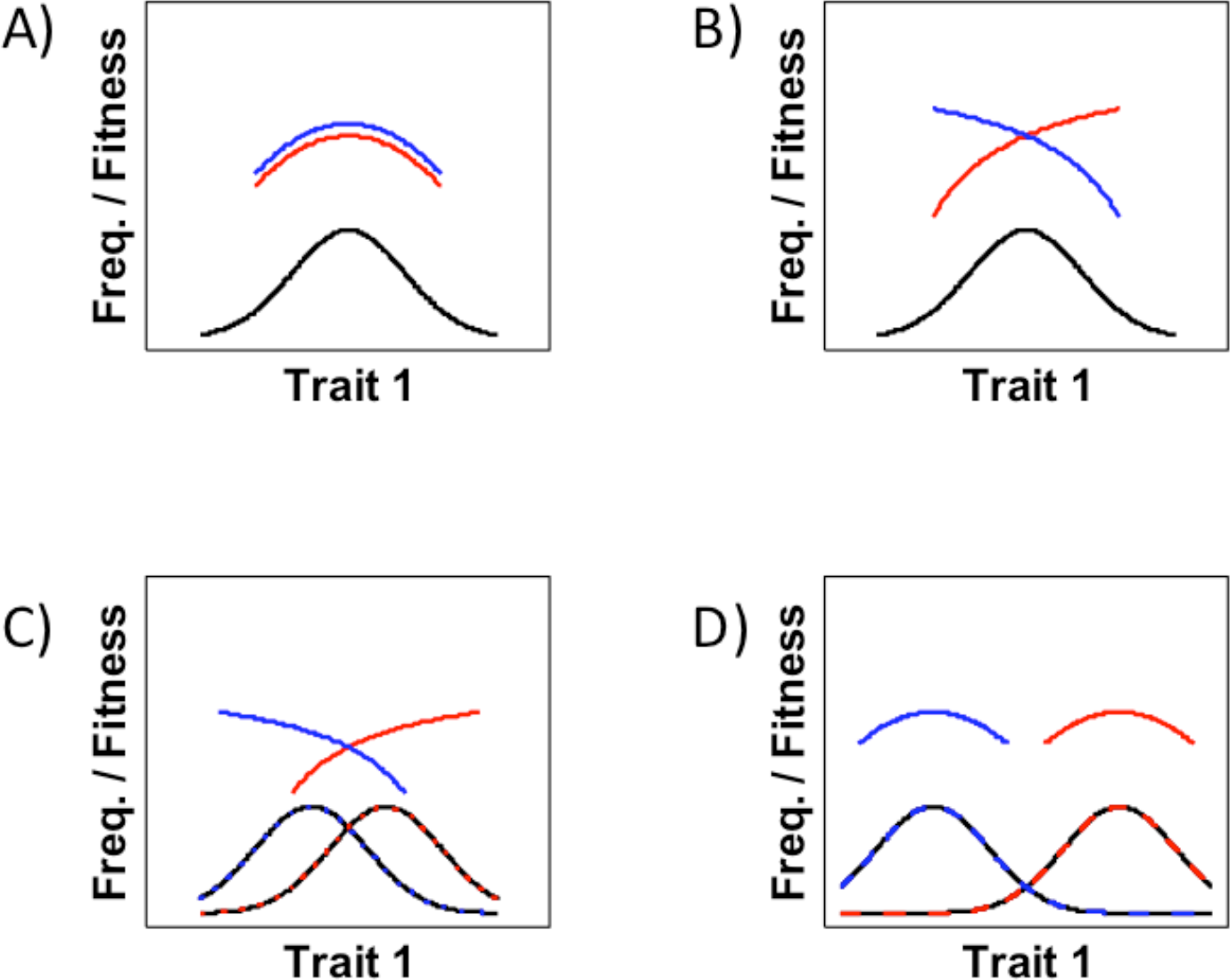
Predicted stages in the resolution of sexual antagonism. A. Initially, the trait is monomorphic and under weak stabilizing selection. B. A change in the physical or social environment causes the previously concordant trait to become subject to opposite patterns of sex-specific directional selection. C. Sexual dimorphism then evolves, causing the sexes to come closer to their respective phenotypic optima, but some antagonism remains. D. Finally, the sexes achieve their optima and the antagonism is completely resolved. Redrawn after information presented in [16].

Most research to date has focused on sexually dimorphic traits, under the assumption that this dimorphism is an indicator of sex-specific phenotypic optima. However the stage of the most severe antagonism and the largest gender load [45,46] is in fact before the trait in question becomes sexually dimorphic, and gene expression data from *D. melanogaster* suggested that most sex-biased genes had already reached their phenotypic optima [47]. In addition, if sexual antagonism results from correlated expression of different traits across the sexes, monomorphism in a given trait may not be informative about its likelihood of being subject to sexual antagonism [48]. This speaks in favour of casting a broad net when searching for sexually antagonistic loci, and not only investigating traits that are already sexually dimorphic.

Proposed mechanisms for the resolution of sexual antagonism include the evolution of sex-linked modifiers, alternative splicing, or gene duplication [38,49]. Gene duplication is a popular theory as to how genes can escape sexual antagonism, by allowing each copy to evolve independently for each sex [50]. Specifically, this would include genes that are activated by sex hormones or have sex-specific methylation, and are thus expressed at different levels in each sex. There is debate about the time-scale of the resolution of sexual antagonism [49,51–55], but regardless of whether the process is fast or slow in evolutionary time, the outcome is always sex-specific genetic architecture. In this sense, sex-specific genetic architecture in disease is likely to be an indirect result of past sex-specific or sexually antagonistic selection. Sexual antagonism or sex-specificity could contribute towards disease risk directly, in which case the sex-specific or sexually antagonistic alleles are themselves a risk factor [14], or indirectly, because in its resolved state, sexual dimorphism creates its own intrinsic sex-specific risk factors, such as behavioural and anatomical specialisation.

## 5. How sex-specific selection affects disease architecture

One implication of sexual antagonism is the maintenance of deleterious genetic variation at higher population frequency than would be expected from mutation-selection balance [43,44]. This leads us to consider its role in susceptibility to common, genetically complex disorders. Consistent with this reasoning, mathematical simulation predicts that alleles that are under sex-differential selection (including sexually antagonistic ones) contribute disproportionately to genetic variation underlying disease phenotypes [14]. We now discuss in greater depth how sexual antagonism for standing genetic variation might contribute to the genetic architecture of complex traits.

### Unequal endophenotype outcome

For common, complex diseases, risk loci are unlikely to cause disease directly, rather they affect a quantitative trait that is biologically linked to the disease (an endophenotype) and becomes a risk factor when the trait value exceeds a certain threshold [56]. This is derived from the endophenotype hypothesis of psychiatric disorders [57], but can be extended to many disorders, examples of which include the relationship between adiponectin level and Type 2 Diabetes [58], and triglyceride level with coronary artery disease [59].

For a correlated trait (with the same genetic architecture between the sexes) an extreme endophenotype value might be beneficial to one sex but detrimental to the other because of other factors, such as sexual dimorphism in physiology, behaviour or environmental exposure. In practice, the extreme trait value might not be beneficial to either sex, but as long as it remains neutral or weakly deleterious, then the causative alleles persist. One example could be for dyslipidaemia that increases risk of myocardial infarction in men but not in women, likely due to the anti-oxidant effects of oestrogen [60].

### Equal disease risk but with unequal fitness effects

Fitness comprises both survival and reproductive components. One might implicitly assume that the reduction in fitness caused by disease is due to disease-related reduction in survival. However, the effects of disease on the second major component of fitness, reproduction, are not always equal between the sexes. One example of this is schizophrenia, where males have a consistently greater reduction in reproductive success than females [61–63]. A second example is for congenital hypothyroidism, associated with loss in fecundity in women but not in men [64]. These examples illustrate how although the genetic architecture of disease may be the same, the fitness effect on each sex as a result of the disease differs.

### Sex-specific migration

It has been suggested that the genetic variation for a sexually antagonistic trait may vary between populations [44], and thus immigration results in the introduction of novel, sexually antagonistic alleles into the host population. Sex-biased immigration will cause alleles that are beneficial to that sex (and thus under net positive selection) to be rapidly introduced into a host population, only for the opposite sex to inherit novel deleterious alleles.

The same principle could be applied to resolved antagonism. For example, methylation is both sex-specific [65,66] and population-specific [67]. It is also proposed as a means by which sexual antagonism can be resolved because it prevents a deleterious allele from being expressed in one sex. The sex-specific migration results in novel combinations of methylated genes increasing the prevalence of extreme (deleterious) phenotypes.

Although obtaining empirical evidence for these processes may be challenging, there is good evidence for large-scale, sex-specific migrations amongst historical human populations from Central Asia [66,68], the Iberian Penninsula [69], the British Isles [70,71], Central Africa [72], Indonesia [73], and globally [74,75]. Furthermore, these mechanisms could provide a novel explanation for the outbreeding depression observed in wild animal populations.

### Sexually antagonistic pleiotropy

We define sexually antagonistic pleiotropy as the deleterious effect of an allele on a fitness-related phenotype in one sex, with a gain in fitness in the other sex through a different phenotype (Figure 2b-c). One example of this comes from quantitative genetics in which cholesterol levels in men are inversely correlated with height in women [76]. Body height in humans is sexually antagonistic, with high values increasing male fitness but reducing that of females [42]. Thus, selection for shorter females causes a maladaptive response in males by raising their cholesterol levels. Indirect empirical evidence indicates that pleiotropic genes are indeed less able to escape sexual antagonism [47,77], and thus the involvement of pleiotropic genes in disease risk seems likely to be amplified by sex-specific selection.

## 6. Methods for identifying sex-specific genetic architecture

For SNP-based association testing, the basic approach to identifying sex-specific effects is to analyse each sex separately, i.e. sex-stratified. In comparison to joint tests, this approach is limited due to the loss in power caused by partitioning of the sample [78]. A common follow-up to the sex-stratified tests is to determine whether the association statistics for each sex are significantly different from one another. Many main-effect studies incorporate sex as a covariate into the analysis, i.e. they are controlling for the effect. However, whilst this approach acknowledges sex-effects it doesn’t allow for their detection. For binary traits with a prevalence of less than 1% inclusion of known covariates actually reduces power [79].

A joint analysis that incorporates a genotype-by-sex interaction term tests the difference in allele frequencies between male and female cases, given their allele frequencies in controls. It is thus more suited to identifying genetic differences in trait architecture between males and females rather than for main effects. The regression model with which to test for genotype-by-sex interactions in an unrelated population sample, is: Y[G,S]=β_0_+β_G_G+β_S_S+β_GxS_(G×S)+ε, where Y is the phenotype value, G is the genotype, S is the sex, β is the standardised regression coefficient of each variable, and ε is the error [80]. Other covariates, such as those used to correct for population stratification, can also be incorporated into this model. The tests can be performed using open-source software, e.g. PLINK [81] and GenAbel [82], although to the best of our knowledge only one GWAS [83] and two (candidate gene) studies have done so [84,85]. For family trio data, a joint, interaction analysis is also possible, exemplified by use of a custom-designed case/pseudo-control test that detected two loci for autism risk [86]. Meta-analysis of GWAS data is a powerful and routine approach to increase power in large heterogeneous sample collections, and an algorithm has been developed in which both sex-specific and main effects can be tested for in a meta-analysis without loss in power [87,88]. An alternative approach (developed for GxE or GxG interactions but applicable to GxS) is to use only loci for which the variance differs significantly between genotypes [89]. This reduces the number of tests whilst enriching for loci most likely to have an interaction component.

The statistical behaviour of genotype-by-sex tests must be assumed to be similar to genotype-by-environment tests, in which the interaction term is binary, obligatory and ideally has the same distribution in case and control populations. Power calculations can potentially be undertaken using such software as Gene-Environment iNteraction Simulator (GENS) [90] and GxEscan [91]. Analytical hazards when using an interaction term include artifactual population substructure [92] and incorrect control of confounders [93] other than sex, such as age, ethnicity, or socio-economic background. Interesting opposite effect direction effects may arise [24,94] but are hard to interpret without replication or biological validation.

As more sex-specific analyses of GWAS datasets are performed, it would be informative for authors to present sex-specific values for i) Trait heritability, ii) The phenotypic variance accounted for by significant SNPs, iii) Genomic prediction/Risk profile score. Finally, given the extent of sexually dimorphic interaction networks [4,95,96], pathway enrichment and epistasis testing should be rewarding.

## 7. Conclusions

Despite sharing genetic variation, there are profound biological differences between males and females. This can result in different optima for shared traits, sexual antagonism, and sexual dimorphism. Sex-specific selection on an allele can have important effects on its maintenance within a population, potentially allowing deleterious, disease-associated alleles to persist [14,43,44]. This predicts the existence of sex-specific architecture, and indeed the recent analyses of large GWAS data sets has brought about an unprecedented rise in the number of robust sex-specific effects in traits of medical relevance (thirty-four loci for twenty-two traits). In fact, we are aware of only one ‘sex-sensitive’ GWAS that did not reach genome-wide significance [83]. Sex differences in disease presentation are often stated as the reason for investigating sex-specific genetic effects, but given that sexual dimorphism is a resolution of sexual antagonism, sexually monomorphic traits are more likely to harbour unresolved conflict and thus also have sex-specific genetic architecture.

Although we have partially excluded sex hormones from sex differences in genetic architecture, they are a major driver of sex-specific gene expression [4]. Furthermore, as their mode of action is to activate transcription factors, SNPs that alter binding-sites for sex hormone-induced transcription factors will have a sex-dependent effect on gene expression. This mechanism is likely to explain some of the sex-specific GWAS results. Alternatively sex-specific epigenetic modification e.g. methylation [97], will inhibit gene expression, masking any functional variation in one sex but not the other. One example of this is known for the ZPBP2 gene and asthma [98].

Much of what is known about sexual antagonism has been obtained through studies on wild and laboratory animal populations, as well as mathematical modelling. Identification of the molecular genetic basis of fitness and of sexual antagonism in model organisms would not only confirm the empirical observations but also provide a grounding for studies of sex-specific genetic architecture in humans. Equally so, ecological studies in humans could also provide interesting perspectives, for example how ecological factors influence selection on specific traits to produce varying degrees of sexually concordant or sexually antagonistic selection across populations [99]. There is evidence from divergent species of weak sex-specific *trans*-eQTLs [26,100], sex-specific residual genetic architecture [101] and sex-specific epistasis [33]. These studies indicate that many modifier loci for common, complex disorders could be sex-specific. One example of this is age-at-onset of Parkinson’s disease being reduced in males only, by the catechol-O-methyltranferase gene Val158 allele [102]. Many monogenic disorders originate from mutations in sex-linked or mitochondrial genes that, because of their transmission dynamics, are under sex-dependent selection. However, the role of common genetic variation on these chromosomes in complex traits is limited by lack of coverage of genotyping chips and suitable analytical methods.

We anticipate that analysis of GWAS data with respect to sex, encouraged by both evolutionary genetics and recent results presented in this review, will generate many more significant findings and highlight the potential role of sex-specific and sexually antagonistic selection as a potent force in human genetic architecture. Finally, we hope that the identification of sex-specific genetic aetiologies in what otherwise appears to be the same disease will result in the development of more effective, sex-specific therapies.

## Acknowledgements

The authors wish to thank laboratory colleagues – Dr Fiona Ingleby, Ms Tanya Pennell, Ms Katrine Lund-Hansen – for their careful reading of the manuscript and insightful comments.

